# PhyteByte: Identification of foods containing compounds with specific pharmacological properties

**DOI:** 10.1101/2020.01.10.902197

**Authors:** Kenneth Westerman, Sean Harrington, Jose M Ordovas, Laurence D Parnell

**Affiliations:** Nutrition and Genomics Laboratory, JM-USDA Human Nutrition Research Center on Aging at Tufts University, Boston, MA USA; Clinical and Translational Epidemiology Unit, Massachusetts General Hospital, Boston, MA USA; Notemeal, Inc., Boston, MA USA; USDA Agricultural Research Service, Nutrition and Genomics Laboratory, JM-USDA Human Nutrition Research Center on Aging at Tufts University, Boston, MA USA

**Keywords:** Bioactivity, Food, Molecule, Natural compound, Nutrition, Protein target

## Abstract

**Background:** Phytochemicals and other molecules in foods elicit positive health benefits, often by poorly established or unknown mechanisms. While there is a wealth of data on the biological and biophysical properties of drugs and therapeutic compounds, there is a notable lack of similar data for compounds commonly present in food. Computational methods for high-throughput identification of food compounds with specific biological effects, especially when accompanied by relevant food composition data, could enable more effective and more personalized dietary planning. We have created a machine learning-based tool (PhyteByte) to leverage existing pharmacological data to predict bioactivity across a comprehensive molecular database of foods and food compounds.

**Results:** PhyteByte uses a cheminformatic approach to structure-based activity prediction and applies it to uncover the putative bioactivity of food compounds. The tool takes an input protein target and develops a random forest classifier to predict the effect of an input molecule based on its molecular fingerprint, using structure and activity data available from the ChEMBL database. It then predicts the relevant bioactivity of a library of food compounds with known molecular structures from the FooDB database. The output is a list of food compounds with high confidence of eliciting relevant biological effects, along with their source foods and associated quantities in those foods, where available. Applying PhyteByte to the *PPARG* gene, we identified irigenin, sesamin, fargesin, and delta-sanshool as putative agonists of PPARG, along with previously identified agonists of this important metabolic regulator.

**Conclusions:** PhyteByte identifies food-based compounds that are predicted to interact with specific protein targets. The identified relationships can be used to prioritize food compounds for experimental or epidemiological follow-up and can contribute to the rapid development of precision approaches to new nutraceuticals as well as personalized dietary planning.

## Background

While a select set of essential nutrients for humans has been well characterized, there is an abundance of lesser-known compounds in the human diet, representing a type of exposure that has been referred to as the “dark matter” of the human exposome [1–2]. These dietary bioactive compounds can have meaningful effects on human phenotypes, to the extent that some, such as lutein and several flavonoids, are under discussion for the establishment of dietary recommended intakes [3]. Despite the potentially important cumulative effects of these compounds, little is known about their bioactivity in the body due to the difficulty of experimentally assaying thousands of compounds for activity against thousands of potential gene products, combined with the complexities of absorption, microbial interactions, and metabolism [4]. Cheminformatic methods, including quantitative structure activity relationship (QSAR) models, can provide *in silico* approaches to prioritize compounds and foods in experimental and epidemiological settings when only the structure of a food compound is known. Pharmaceutical drugs can provide a critical set of anchors for such models, as their primary biological mechanisms of action are typically well characterized.

Computational approaches to generating hypotheses related to food and food compound bioactivity have been introduced [5–6]. However, existing methods have focused primarily on literature mining based on natural language processing, rather than optimizing for the output of food compound activities related to a given input gene or protein of interest. Methods described to date have used relatively basic QSAR methods, such as comparisons based on Tanimoto similarity scores, which may fail to capture important signals. Additionally, there can be significant utility in identifying the food(s) that contains a compound of interest both as a source material or in the formulation of a novel product. The growth of relevant databases containing pharmaceutical and food composition information continually offers opportunities to revisit and improve QSAR tools. The United States Department of Agriculture (USDA) has a long history of producing high-quality data for its food composition databases [7], and inclusion of established or potential health effects would be a useful extension of these data.

Here, we develop and demonstrate a machine learning-based approach, PhyteByte, that assigns putative bioactivity to food compounds based on a training set of pharmaceutical drugs. We show the efficacy of PhyteByte using the specific example of PPARG, the known target of the thiazolidinedione (TZD) drug class.

## Implementation

In order to identify functional relationships between a food compound and a drug, along with its associated bioactivity data, we used data from two sources: ChEMBL and FooDB. ChEMBL is a manually curated database of almost 2 million (1,879,206 in version 25) bioactive molecules with drug-like properties [8–9]. These data were retrieved from ebi.ac.uk/chembl/ on 9/27/2019. FooDB (version 1.0) is a comprehensive resource on food constituents, chemistry and biology, with over 85,000 compounds in its repository [10]. These data were accessed from foodb.ca on 9/27/2019. As allele-specific binding data are not available in ChEMBL, PhyteByte currently does not have the means to incorporate genetic variants into its prediction.

The PhyteByte computational pipeline is outlined in Figure 1 (along with details related to a specific gene input; see **Results & Discussion**). The processing of data through PhyteByte is initiated by selection of an input protein target query, from which drugs acting on that target (sourced from ChEMBL) are obtained to provide computational fingerprints of their molecular structure. The fingerprints are processed by a predictive model to yield likely bioactivity for food compounds (sourced from FooDB), which in turn are queried in FooDB to retrieve foods containing those compounds, with quantified amounts where available.

**Figure 1.**
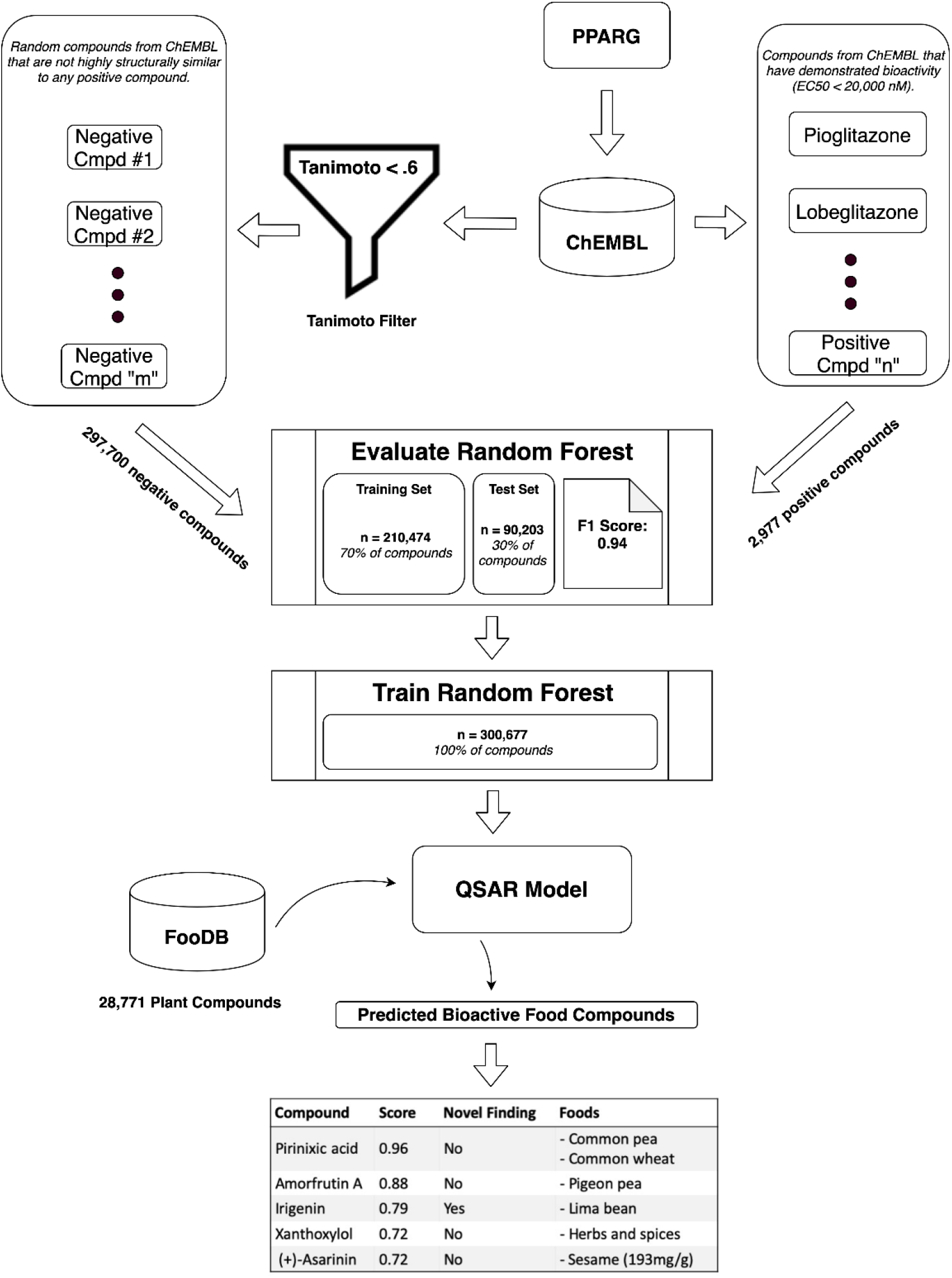
Schematic data flow for PhyteByte from protein target input to predicted bioactive food compounds.

Specifically, a target specification (provided in the form of an HGNC gene symbol) serves as input for a query to ChEMBL that retrieves chemical structures for molecules with evidence of relevant bioactivity for the protein encoded by that gene. Bioactivity is defined as an IC50 (inhibitory concentration: the concentration of the molecule required to inhibit the biochemical function of the target by 50%) or EC50 (effective concentration; the concentration of the molecule required to induce 50% of the maximal response or effect on the target) of <20,000 nM based on the user-specified compound effect type (antagonist vs. agonist). Because ChEMBL does not contain explicit annotations as to the effect type, a heuristic is used in which the strength of antagonists and agonists are evaluated using IC50 and EC50 values, respectively. Compound structures are retrieved as simplified molecular-input line-entry system (SMILES) strings. SMILES strings are a dense, character-based representation of chemical compounds (for example, “COC1=CC(=CC(=C1OC)O)C2=COC3=C(C2=O)C(=C(C(=C3)O)OC)O” for irigenin, a compound in Table 1). The SMILES strings are then converted into FP2 binary fingerprints using the Pybel Python package [11], which acts as a wrapper for the OpenBabel chemical file format interconversion tool. FP2 fingerprints are a binary compound representation (as a 1024-bit vector) formulated based on the occurrence of specific linear fragments up to 7 atoms in length. Further details on the SMILES and FP2 formats are available from the Open Babel publication [12] and online Wiki (https://openbabel.org). A set of negative examples, chosen to be 10 times the size of the positive set, is also retrieved at random from the full set of ChEMBL molecules. The negative examples are converted to FP2 fingerprints after filtering such that no negative compound has a Tanimoto similarity score >0.6 with any molecule in the positive set. The Tanimoto coefficient is defined as an association coefficient (in comparison to a distance coefficient) that measures similarity, here as chemical similarity based on SMILES representation of the molecule [13]; formulae for the Tanimoto coefficient are presented elsewhere [14]. No explicit upper limit for molecular mass of the bioactive molecules is set, but we note that the vast majority (>98%) of molecules in ChEMBL are categorized as small molecules.

**Table 1.**
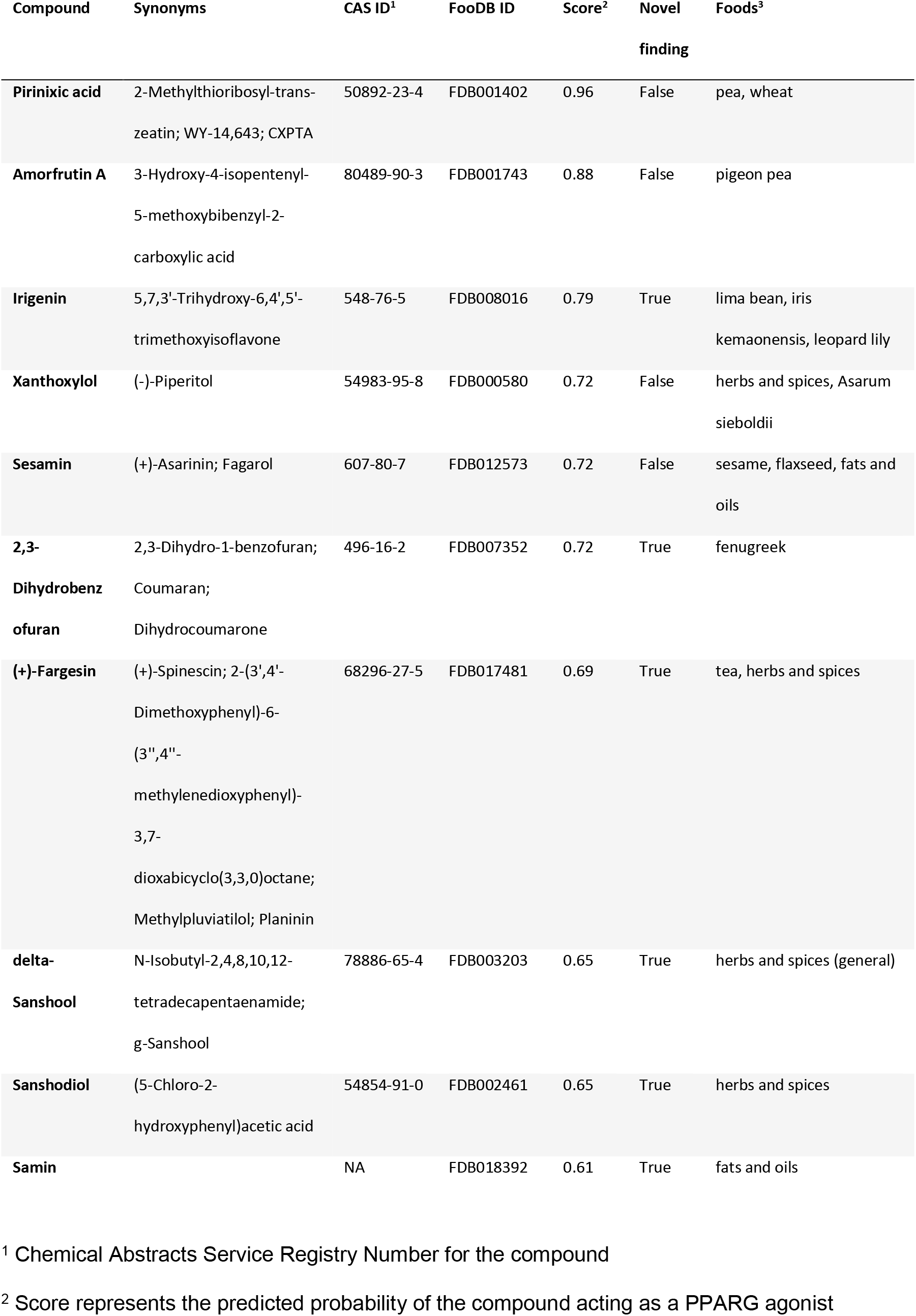
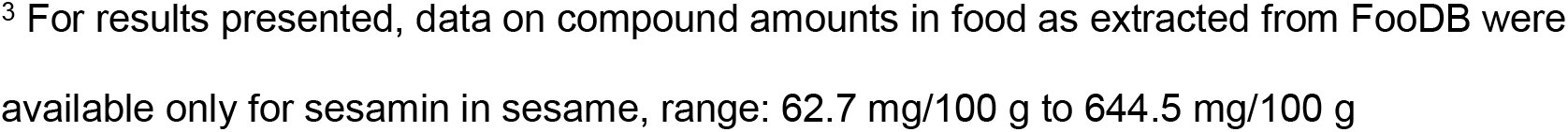
Top food compound results from PhyteByte for input of PPARG.

Next, a random forest model is trained (using the sklearn Python package) to classify compounds as to their bioactivity against the protein of interest. Inputs consist of the binary fingerprints (a binary feature vector of length 1024) and class labels (positive if evidence of bioactivity for the target exists in ChEMBL, or negative if not). The random forest classifier is an ensemble learning method that trains a set of independent decision trees to discriminate between positive and negative examples. Given a new compound (in this case, a food compound), binary predictions from each individual decision tree are averaged to output a probability of bioactivity. Models in PhyteByte use 100 component trees, with all additional parameters following sklearn defaults. The training and testing dataset split is created by assigning a random 30% of compounds to the testing dataset (including a consistent random seed for reproducibility), with the remaining 70% assigned to the training dataset. We note that after evaluation, the final model used to process food compounds is trained on the full dataset. An initial indication of model performance is evaluated in a 30% held-out testing set using the F1 score, or the harmonic mean of precision and recall. This metric is calculated as 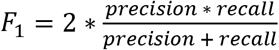 where precision is the fraction of predicted bioactive compounds that have evidence for bioactivity in ChEMBL, and recall is the fraction of compounds with evidence for bioactivity in ChEMBL that are predicted to be bioactive. True positive (TP) is defined as bioactivity in ChEMBL and predicted to be bioactive; false positive (FP) is defined as no bioactivity in ChEMBL but predicted to bioactive; false negative (FN) is defined as bioactivity in ChEMBL but not predicted to be bioactive. Thus, precision = TP / (TP + FP), recall = TP / (TP + FN), and F1 is calculated as above.

Using this trained model, the full set of food compounds available from FooDB are then characterized as to their probability of bioactivity with respect to the input protein. The list of probable dietary bioactive compounds is presented as output, along with their concentrations in foods as available in FooDB and an indication of whether the relationship is novel (i.e. does the compound lack existing evidence of bioactivity for the input protein in ChEMBL?). PhyteByte source code and installation instructions are available at https://github.com/seanharr11/phytebyte, and as a standalone tarball in Additional file 1 (capturing this repository at the time these analyses were performed).

## Results & Discussion

We have demonstrated the functionality and output of PhyteByte using the input gene *PPARG* (CHEMBL235), whose protein product is the target of the thiazolidinedione (TZD) drug class. TZDs are widely prescribed to treat type 2 diabetes, and additionally may have broader cardiometabolic benefits [15]. However, TZDs also have documented side effects and FDA-issued alerts of adverse effects [16], suggesting a potential benefit of identifying alternative or complementary food-based bioactives. Details of the PhyteByte pipeline as realized for PPARG agonists are presented in Figure 1. 2977 positive compounds were retrieved from ChEMBL, along with 297,700 negative compounds. The trained model exhibited an F1 score (harmonic mean of precision and recall) of 0.94 in a 30% held-out set, indicating a reasonably strong discriminative capacity within the set of molecules in ChEMBL. This score may be biased upwards due to limitations in the set of pharmaceutical compounds explored to date, but nonetheless indicates an ability to classify potential food compounds effectively.

When used to score compounds from FooDB, the model identified a series of molecules with potential agonist bioactivity for PPARG. Table 1 lists the 10 molecules with a predicted bioactivity confidence of greater than 0.60 that also had associated foods in FooDB; tabulated results include the identified food compound, common synonyms, CAS and FooDB identifiers, PhyteByte output score, whether the compound-PPARG interaction is a novel finding, and foods reported to contain that compound. Molecules such as pirinixic acid (or WY-14643) and xanthoxylol have been shown to activate PPARG [17–19], albeit the latter only as an activator of *PPARG* transcription [20]. Other molecules have little to no existing evidence in the scientific literature of acting as PPARG agonists. These include irigenin (an O-methylated flavone found in lima bean), sesamin (a lignan found in sesame and flaxseed), fargesin (a lignan from tea, herbs and spices), delta-sanshool (an n-acyl amine from herbs and spices), and the lignan sanshodiol (from herbs and spices). Such molecules could be prioritized for detailed experimental validation. Complete output of PhyteByte for PPARG as input and resulting identified compounds scoring above 0.50 is presented in Additional file 2.

Tools such as PhyteByte consider only small molecules and are limited by the content of the input databases. Importantly, these resources are expected to become increasingly comprehensive, especially for food compounds. For example, efforts are underway by the USDA to expand their food composition databases [7], and recent investigations have identified additional compounds produced during food processing [21] and by human microbiota [22], which may promote certain health effects. While QSAR models are susceptible to false positives due to activity cliffs (key discontinuities in the structure-activity landscape), outputs from PhyteByte are intended to be only putative structure-activity relationships to be explored further through complementary computational and laboratory methods [23]. Experimental and/or epidemiological assessment eventually will be required to validate at least some subset of the algorithmic predictions before this tool could be used in clinical settings or for dietary recommendations.

In future versions of the software, we anticipate more flexibility in both the inputs and databases. For example, inputs may include phenotypes (to be linked to a set of target gene products and user-defined food compound datasets following a pre-defined schema may be used to complement FooDB. Additionally, as more follow-up testing of food compound-target interactions is performed, those results can be used as a complementary source of interactions for PhyteByte and form the basis for a catalog of all such interactions for a single food. Complementary data streams, such as those based on text mining [5], pharmacology networks [24] or drug interaction data (to identify potential similar food compound interaction effects), could provide additional support for food compound-phenotype links. Future work also should include more fine-grained annotations of positive training molecules (based on type of effect on the target, strength, and mechanism of action) as well as alternative QSAR modeling approaches [25].

## Conclusions

PhyteByte is a machine learning-based tool for discovery of interactions between food compounds and specific proteins or phenotypes. The software enables prioritization of these compounds for future research and hypothesis generation for condition-specific dietary interventions. Applied to the *PPARG* gene, this tool recovered known ligands and generated the basis for new hypotheses useful for cell-based assays or epidemiological inquiries. Our work provides additional proof-of-concept for the emerging field of “computational nutrition” based on food compounds, building on previous research that applied cheminformatic approaches to assign putative biological function to molecules of interest.

## Availability and requirements

Project name: Phytebyte

Project home page: https://github.com/seanharr11/phytebyte

Operating system(s): Unix-based (MacOS, Linux)

Programming language: Python

Other requirements: Python 3.6 or higher

License: AGPLv3

Any restrictions to use by non-academics: License needed

## Abbreviations

EC50: effective concentration
IC50: inhibitory concentration
PPARG: peroxisome proliferator activated receptor gamma
QSAR: quantitative structure activity relationship
SMILES: simplified molecular-input line-entry system
TZD: thiazolidinedione
USDA: United States Department of Agriculture

## Declarations

Ethics approval and consent to participate – not applicable

Consent for publication – not applicable

### Availability of data and materials

All data generated during this study are included in this published article and its supplementary information files. The ChEMBL and FooDB datasets analyzed during the current study are available are available at ebi.ac.uk/chembl/ and foodb.ca [8–10].

### Authors’ contributions

KW, SH and LDP conceived of, designed, wrote and tested PhyteByte, and/or analyzed results. KW, SH and LDP wrote, and all authors reviewed and approved the submitted manuscript. JMO provided financial support to KW and LDP.

## Acknowledgements

Mention of trade names or commercial products in this publication is solely for the purpose of providing specific information and does not imply recommendation or endorsement by the U.S. Department of Agriculture. The USDA is an equal opportunity provider and employer.

## Additional file 2

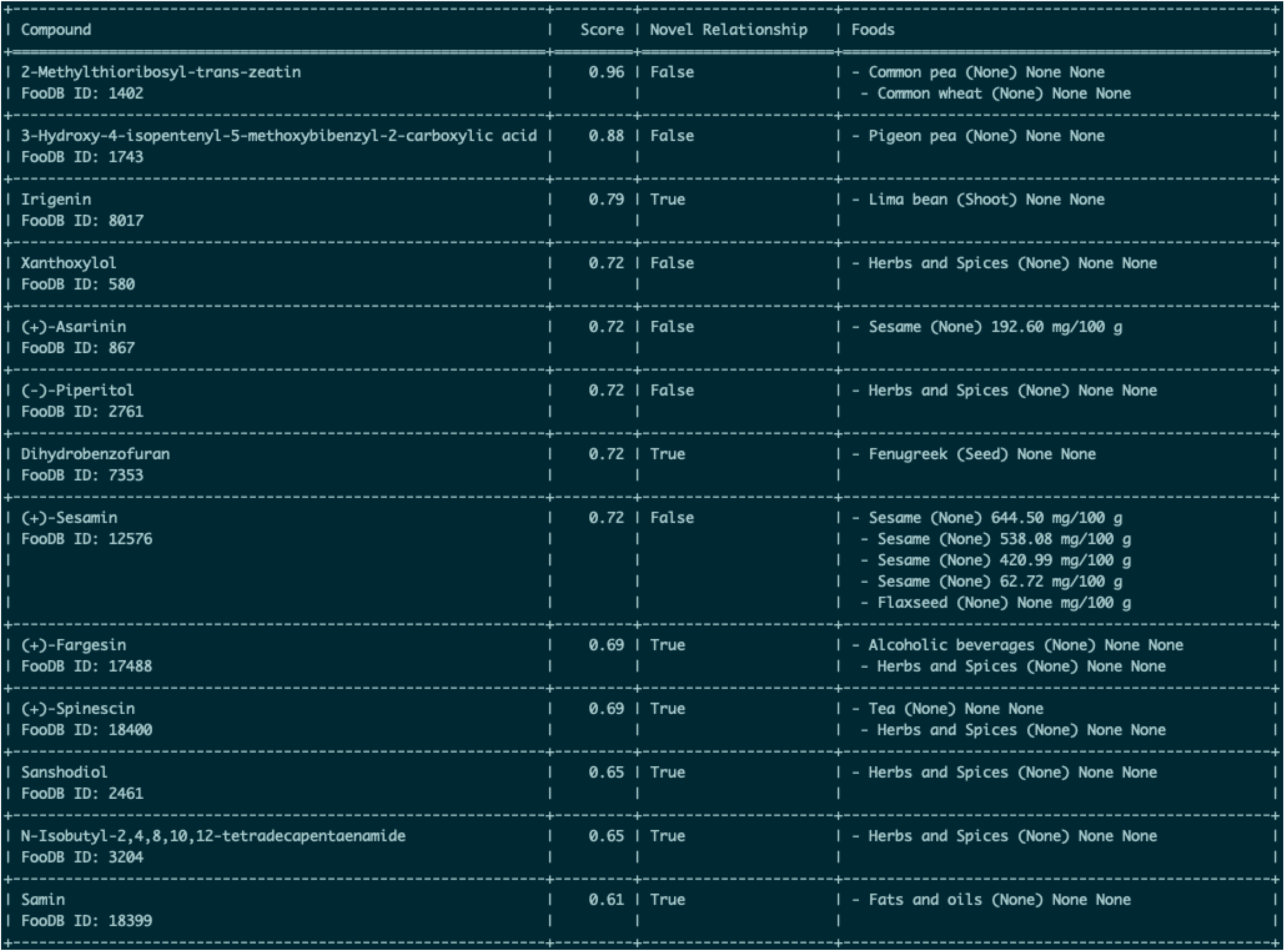

